## Supplemental Tables and Figures for "Arginine Methylation Regulates Ribosome CAR Function"

### 1. Supplementary Materials

#### *1.1 Supplementary Figures*

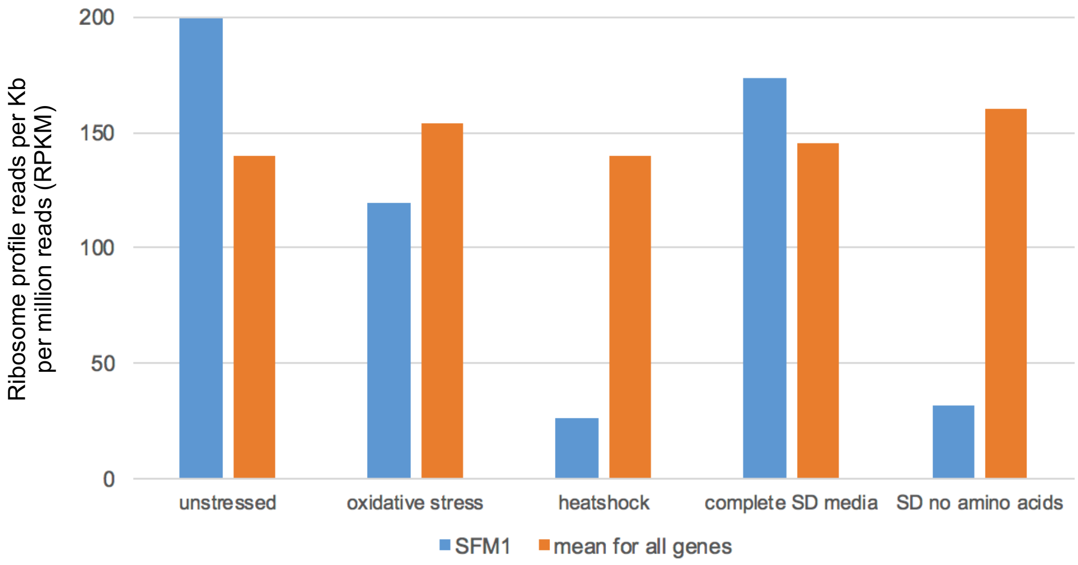


**Figure S1.** *SFM1* expression is depressed in stress conditions.
Analysis of ribosome profile data of Gerashchenko and Gladyshev [21] revealed that Sfm1, the methyl transferase that methylates Rps3 R146, has reduced expression in stress conditions.

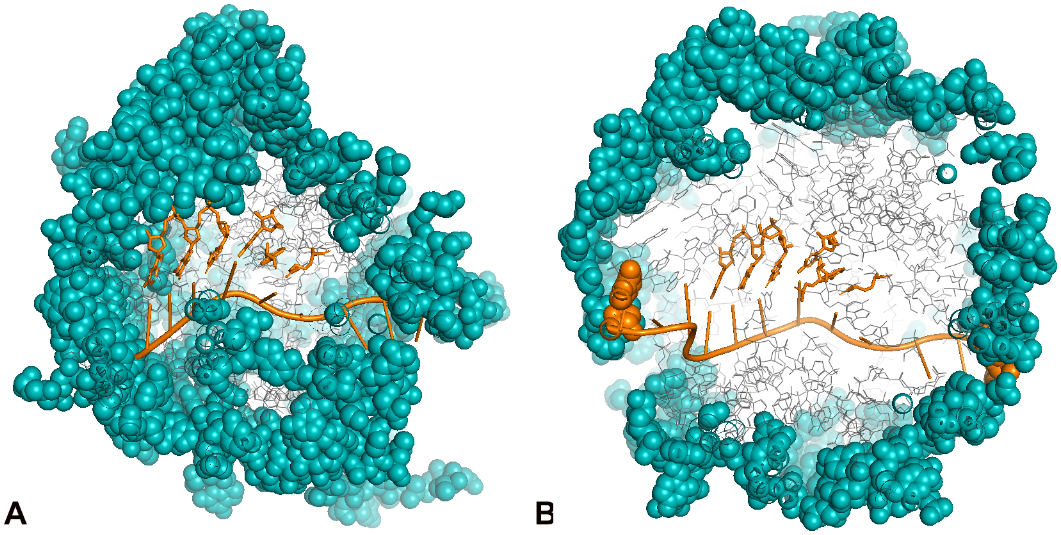


**Figure S2.** Design of N1 and N2 subsystem neighborhoods.
Subsystem N1 was used previously [12] in analysis of the CAR interface. Subsystem N2, used in this study, is more centered over the A site and CAR interface, and exhibits similar behavior to N1. $Na^+$ ions were used to achieve electro-neutrality in N1, and $K^+$ ions were used for N2.

## A. R146 60 ns

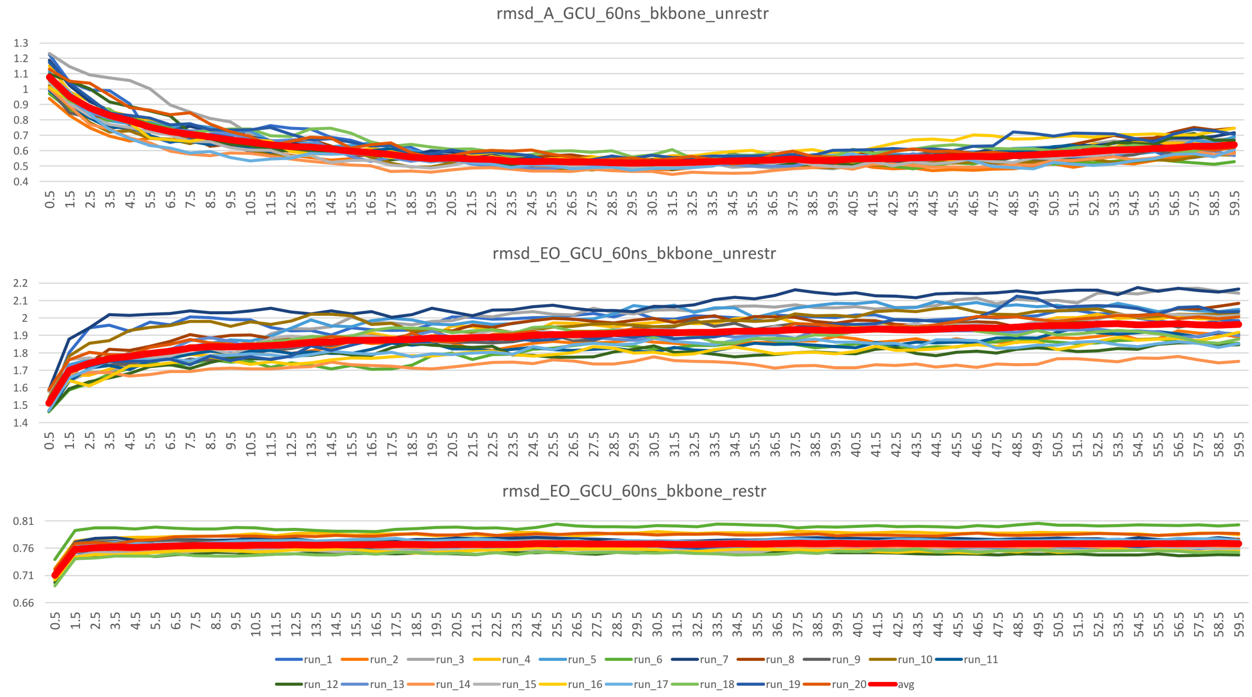


## B. R146 100 ns

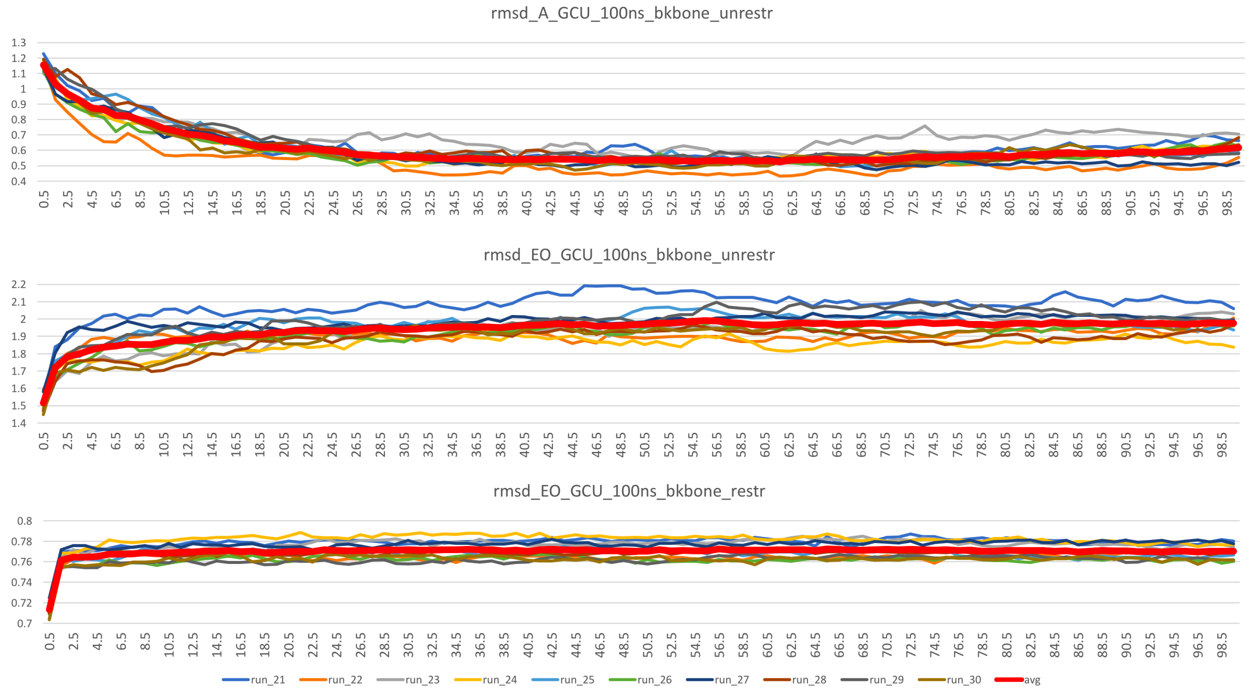

C. mR146 60 ns

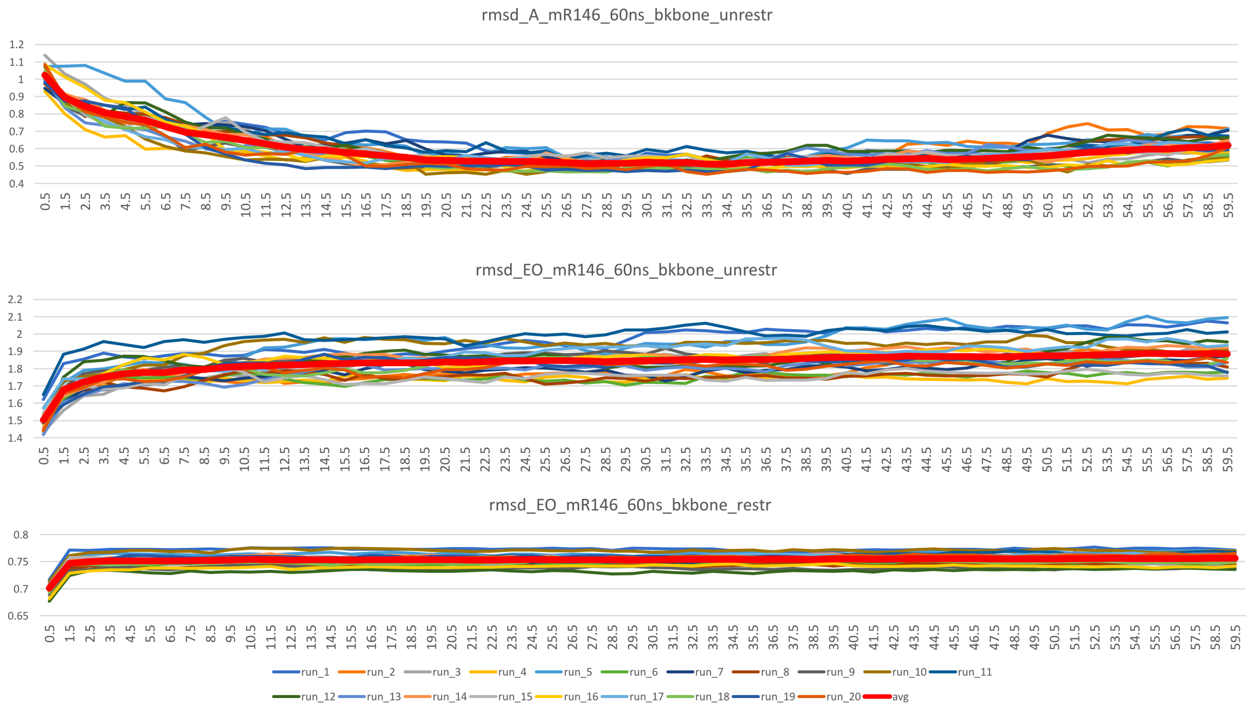


D. mR146 100 ns

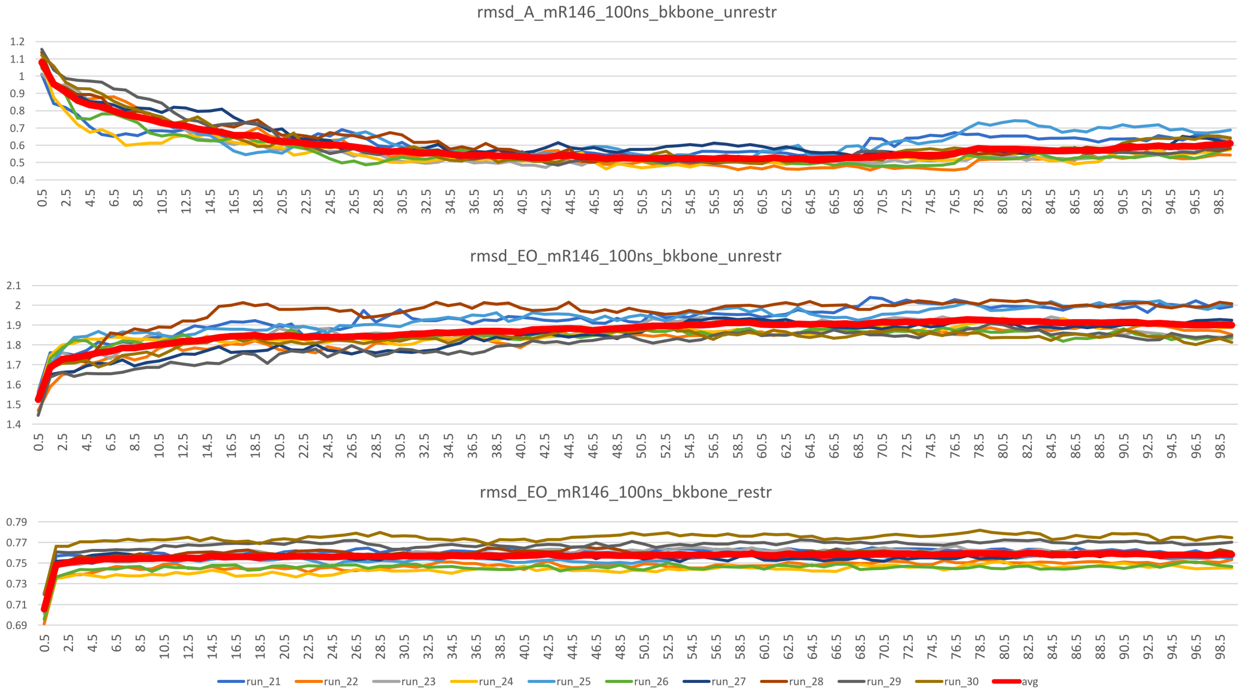


**Figure S3.** RMSD profiles for R146 and mR146 in neighborhood N2.
RMSD profiles were computed using as reference the average structure across the full trajectory (RMSD_A) or the structure at the end of equilibration (RMSD_EO). R146 and mR146 subsystems were analyzed using 60- and 100-ns trajectories. The trajectories stabilized by 20 ns and the first 20 ns of trajectories were not used for further analysis.

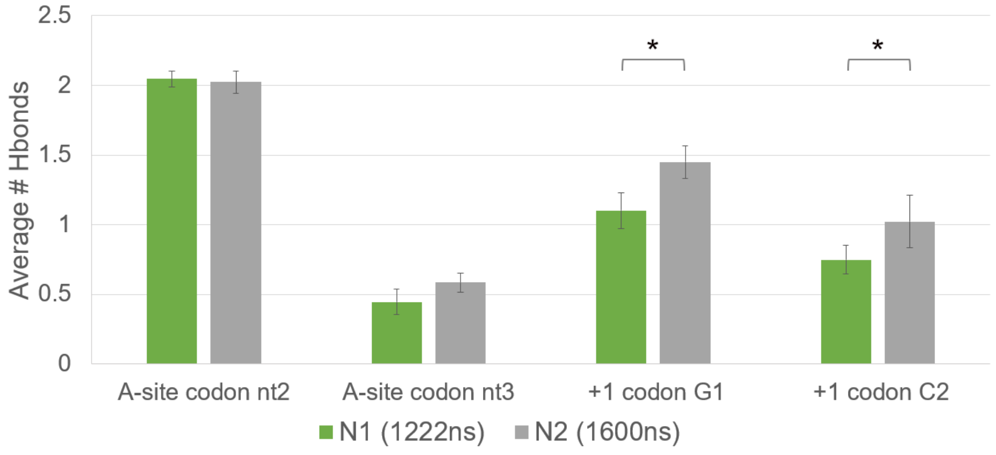


**Figure S4.** CAR H-bonding is stronger in neighborhood N2.
Subsystems N1 and N2 were compared using +1 codon GCU. N2 had greater H-bonding between CAR and G1 or C2 of the +1 codon ($p < 0.05$ *).

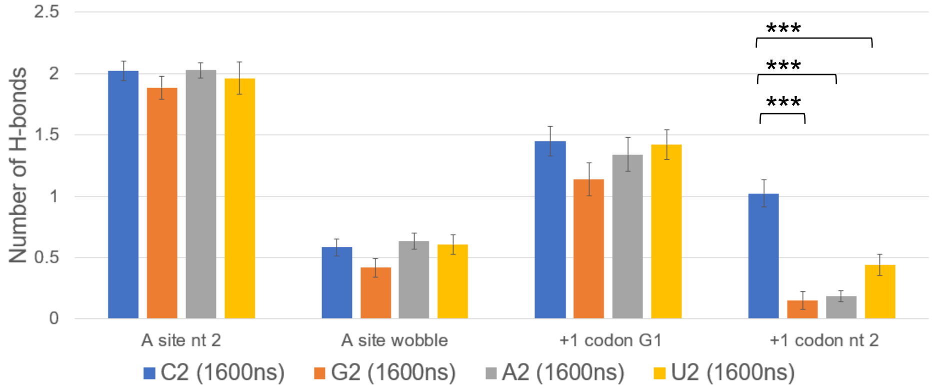


**Figure S5.** C2 substitutions in the +1 codon reduce CAR H-bonding.
As observed previously with subsystem N1 (ref), the N2 subsystem also revealed that H-bonding is reduced when the second nucleotide of the +1 codon (C2) is replaced with G2, A2 or U2 ($p < 0.001$ ***), suggesting that N1 and N2 behaviors are similar.

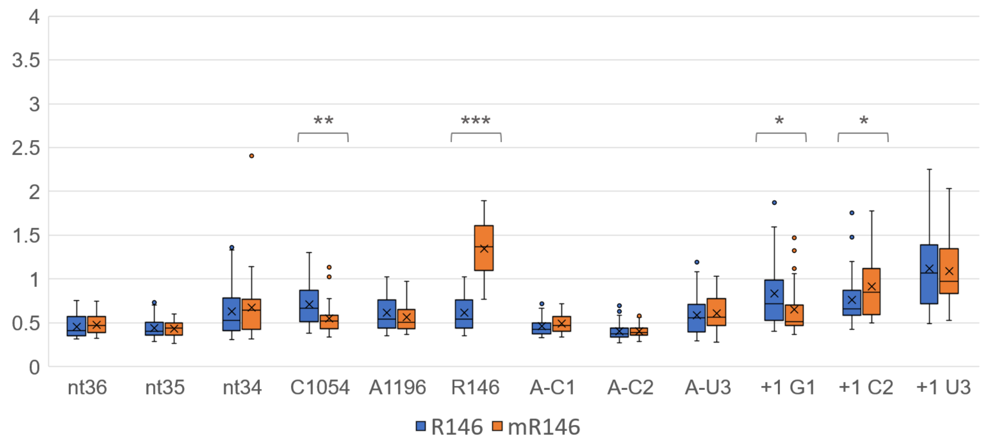


**Figure S6.** mR146 has elevated RMSF
Like the N2 subsystem (Figure 2), the N1 subsystem showed that mR146 has elevated RMSF compared to R146 for core base heavy atoms of nucleotides and guanidinium group heavy atoms ($p < 0.001$ ***).

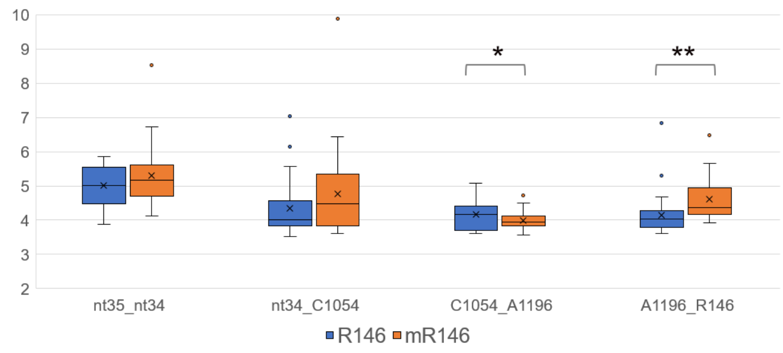


**Figure S7.** mR146 has reduced stacking.
Like the N2 subsystem (Figure 3A), the N1 subsystem showed that mR146 has reduced pi stacking compared to R146 ($p < 0.01$ **), indicated by increased distance between centers of mass of the A1196 base and the guanidinium group.

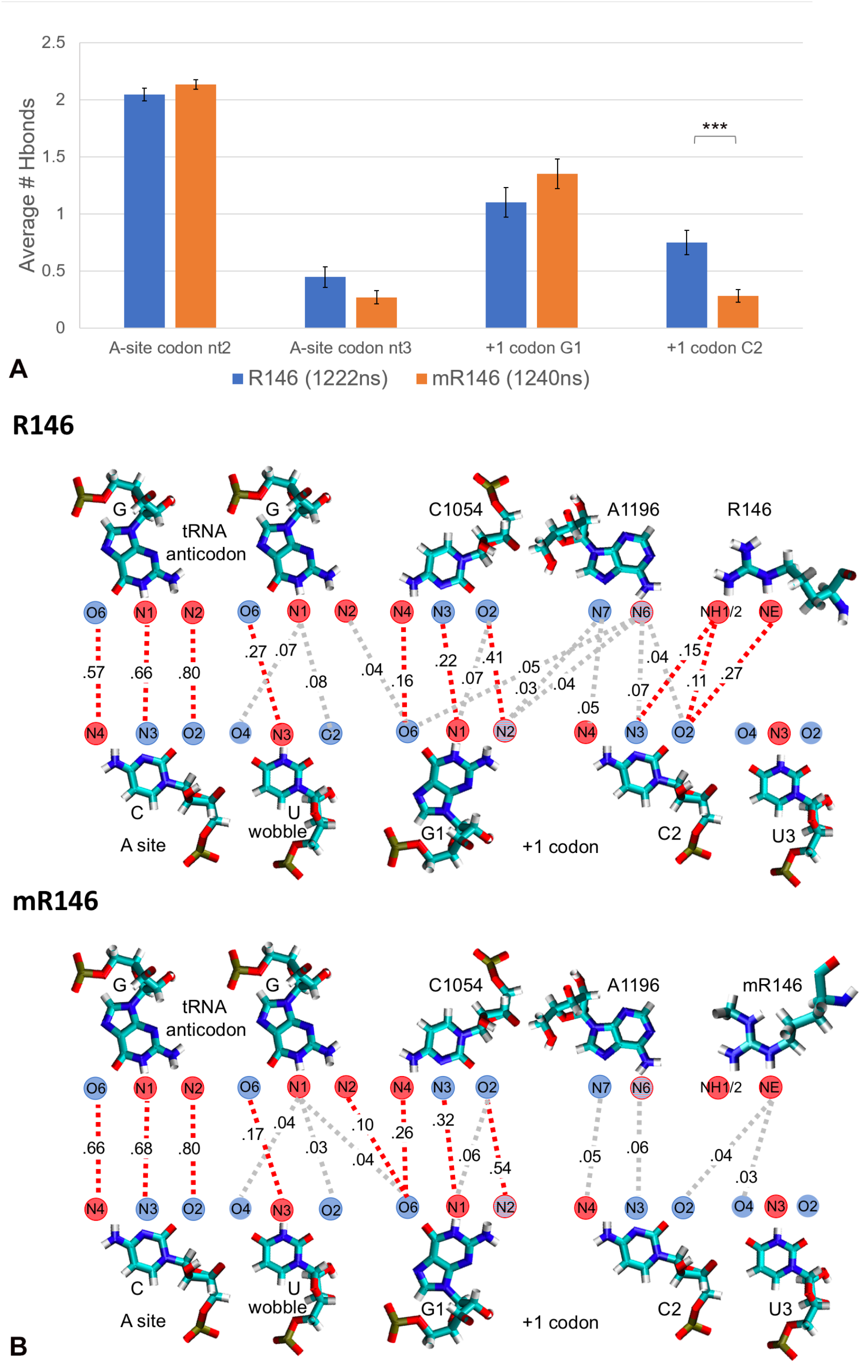


**Figure S8.** mR146 has reduced CAR/+1 codon H-bonding.
(A) Like the N2 subsystem (Figure 4), the N1 subsystem showed that mR146 had reduced H-bonding between the CAR interface and the +1 codon. +1 codon C2 had significantly reduced H-bonding ($p < 0.001$ ***). (B) Details of H-bonding interactions.

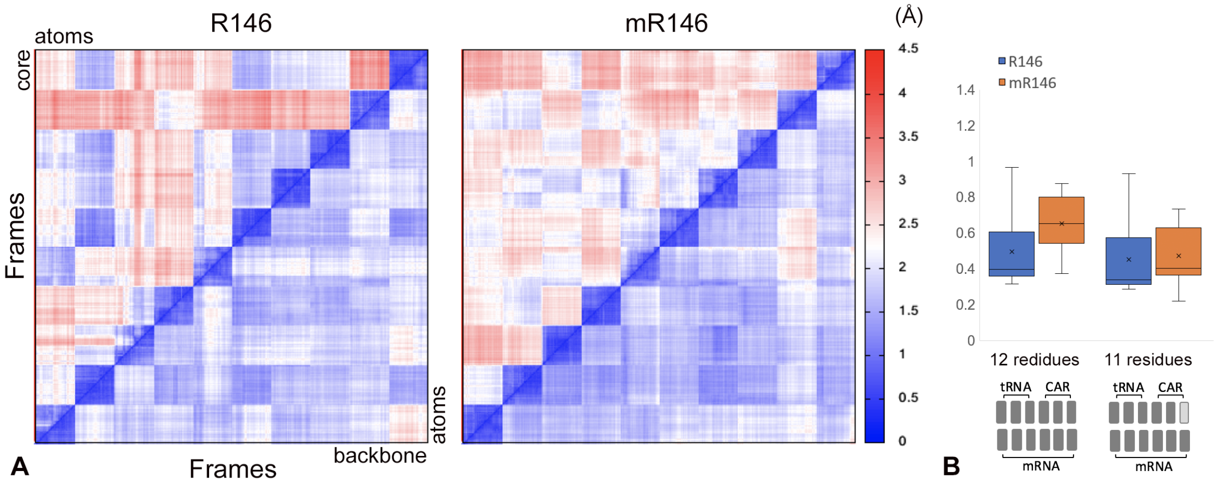


**Figure S9.** RMS2D analysis.
(A) RMS2D analysis was performed for 10 trajectories of 100 ns (excluding the first 20 ns) sampled at 1 frame/ns using backbone atoms (top right triangles) or "core" (base and guanidinium group) heavy atoms (bottom left triangles) of 12 residues (see Figure 5). (B) Similar to analysis of 60-ns trajectories (Figure 5), the differences across experiments comparing backbone and core RMSD were more pronounced for mR146 than R146 (but not significant in bootstrap tests).

*1.2 Supplementary Tables*

**Table S1A.** Unmethylated R146 Trajectories

| R146 Trajectory Number | nt34-CAR Anchoring [1,2,3] | CAR Stacking [4] | CAR / mRNA Stacking [5] | A-site nt3 Interactions | +1 nt2 Orientation [6] | Other Comments |
|---|---|---|---|---|---|---|
| **1** | ✓ | CAR | R to +1 U3 | X | out | |
| **2** | X | CA | A to +1 C2 | nt34 | up | R behind A |
| **3** | ✓* | CAR | R to +1 U3 | C1054 | up | *nt34-C unstacks |
| **4** | ✓ | CAR | R to +1 U3 | nt34 | out | A-site nt1 pointed away from nt36 |
| **5** | X | CAR | X | nt34 | out | |
| **6** | ✓ | CA | A to +1 C2<br>R to +1 U3 | nt34 | out | |
| **7** | (X) | CAR | R to +1 C2 | C1054 | out | U3 pointed down |
| **8** | ✓ | CAR | X | C1054 | up | R-U3 H-bonding |
| **9** | X | X | X | X | out | R behind A,<br>U3 pointed down |
| **10** | X | CAR | R to +1 U3 | X | (up) | |
| **11** | (✓) | CAR | R to +1 U3 | nt34 | up | |
| **12** | (X) | X | A to +1 C2 | nt34 | out | A-site nt1 pointed away from nt36,<br>R behind A |
| **13** | ✓ | CAR | R to +1 U3 | nt34 | up | |
| **14** | X | CA/AR | X | nt34 | up | C1054 perpendicular to nt34 |
| **15** | X | X | A to +1 C2 | nt34 | up | R moves behind A |
| **16** | ✓ | CAR | R to +1 U3 | nt34 | up | |
| **17** | X | CA | A to +1 C2<br>R to +1 U3 | nt34* | (out) | *nt34 rotated 90° |
| **18** | ✓ | CA | A to +1 C2<br>R to +1 U3 | nt34, +1 G1 | out | |

| | | | | | | |
|---|---|---|---|---|---|---|
| **19** | X | CA | A to +1 C2 | nt34 | out | A-site nt1 pointed away from nt36 |
| **20** | X | CAR | R to +1 U3, then R to +1 C2 | nt34 | up then out | |
| **21** | X | CAR* | R to +1 U3* | nt34 | up | *R moves behind A then back to AR stacked |
| **22** | ✓ | CAR | R to +1 U3 | nt34 | up | |
| **23** | ✓ | CAR | X | nt34 | up | A-site nt1 stacked with nt36, U3 pointed down |
| **24** | ✓ | CAR | X | nt34 | up | U3 pointed down |
| **25** | X | CAR | R to +1 U3 | C1054 | out | A-site codon shifts forward 1nt and then back |
| **26** | ✓ | CAR | R to +1 U3 | nt34 | up | |
| **27** | ✓ | CAR/CA | R to +1 U3/ A to +1 C2 | nt34 | out | A-site nt1 pointed away from nt36, A-site nt2 Hbond to nt36 |
| **28** | ✓ | CAR | R to +1 U3 | nt34 | up | |
| **29** | X | CAR | X | nt34 | out | G1 rotated 90°, U3 pointed down |
| **30** | ✓ | CAR | A to +1 C2<br>R to +1 U3 | nt34 | out | |

**Table S1B.** Methylated R146 Trajectories

| mR146 Trajectory Number | nt34-CAR Anchoring[1,2,3] | CAR Stacking[4] | CAR / mRNA Stacking[5] | A-site nt3 Interactions | +1 nt2 Orientation[6] | Other Comments |
|---|---|---|---|---|---|---|
| **1** | ✓ | CAR to CA* | (A to +1 G1) | nt34 | out | *R146 moves behind A1196 |
| **2** | X | CA | (A to +1 C2 and R to +1 U3) | nt34 | out | C1054 rotated 90°, R146 behind A1196 |
| **3** | (✓) | CA | A to +1 C2, | (nt34) | out | R146 behind |

| | | | | | | |
|---|---|---|---|---|---|---|
| | | | (R to +1 U3) | | | A1196 |
| **4** | ✓ | CA(R) | X | nt34 | down | U3 pointed down |
| **5** | X | CA to CAR* | X | C1054 | up | nt34 rotated 90°, *R146 moves from behind A1196 to stacked |
| **6** | ✓ | CAR | R to +1 C2 to +1 U3 | nt34 | out | |
| **7** | ✓ | CA | A to +1 C2 | (nt34) | out | R146 behind A1196 |
| **8** | ✓ | CA | A to +1 C2, (R to +1 U3) | nt34 | out | |
| **9** | X | (AR) | A to +1 G1, R to +1 C2 | X | out | A site nt1 rotated 90° |
| **10** | X | (CR) | (A to +1 C2) | nt34 | out | R146 behind A1196 |
| **11** | ✓ | CA | (A to +1 C2), R to +1 U3 | nt34 | out | A-site codon shifted forward 1nt, nt34 unstacked from nt35 |
| **12** | ✓ | CAR to CA* | (R to +1 C2 and +1 U3) | nt34 | up | *R146 moves behind A1196 |
| **13** | ✓ | CAR | R to +1 U3 | X | up | nt34 rotated 90° |
| **14** | X | CAR | C to +1 G1, (R to U3) | nt34 | up | |
| **15** | (X) | CA(R) | (A to +1 C2) | nt34 | (up) | U3 rotated down |
| **16** | ✓ | CA(R) | (A to +1 C2), (R to +1 U3) | nt34 | (up) | |
| **17** | ✓ to X* | CA to X* | A to +1 G1 | nt34 | out | U3 rotated down, *R146 moves behind A1196 |
| **18** | ✓ | CA | A to +1 C2 | nt34/nt35 | out | A-site codon shifted forward 1nt |
| **19** | ✓ | AR | R to +1 U3 | X | (up) | A-site codon shifted forward 1nt |
| **20** | ✓ | CA | X | X | out | R146 behind A1196 |

| | | | | | | |
|---|---|---|---|---|---|---|
| **21** | ✓ | CAR | R to +1 C2 | X | out | A-site nt3rotated 90° |
| **22** | ✓ | CA | A to +1 C2 | nt35 | out | U3 pointed down |
| **23** | (X) | (CR) | X | nt34 | up | R146 behind A1196 |
| **24** | X | CAR to CA* | X then A to +1 C2 | nt34 | out | G1 pointed toward A-site, *R146 moves behind A1196 |
| **25** | ✓ | CA | X | nt34 | up | R146 behind A1196 |
| **26** | ✓ | CA | A to +1 C2 to +1 U3 | X | up | R146 behind A119 |
| **27** | ✓ | CA | A to +1 C2 | (X) | out | |
| **28** | X | (CR) | (R to +1 C2) | nt34 | out | R146 behind A1196 |
| **29** | ✓ to X | CAR to CA | R to +1 C2 to +1 U3, then X | nt34 | out | |
| **30** | ✓ | CR | X | nt34 | up | R146 behind A1196, G1 stacks with C1054 at the end |

Notes:

[1] Entries in parenthesis show intermittent behaviors.

[2] Entries marked with * are discussed in the Other Comments column.

[3] nt34-CAR Anchoring describes if there was consistent stacking of tRNA nucleotide 34 with C1054 (✓).

[4] CAR Stacking shows reduced stacking with mR146 and the arginine is frequently located behind A1196.

[5] CAR/mRNA Stacking shows cases of CAR residues stacking with +1 codon nucleotides.

[6] +1 nt2 Orientation describes if the 2nd nucleotide of the +1 codon was oriented towards the CAR interface (up).

**Table S2.** Neighborhood 2 (N2) residues.

| Chain | 5JUP Numbering* | 5JUP Restrained | Subsystem N2 Numbering | Subsystem N2 Restrained |
|---|---|---|---|---|
| A 18S rRNA | 7-13 | 7, 12-13 | 1-7 | 1, 6-7 |
| A 18S rRNA | 558-583 | 558-562, 567-572, 582-583 | 8-33 | 8-12, 17-22, 32-33 |
| A 18S rRNA | 1000-1002 | 1000-1002 | 34-36 | 34-36 |
| A 18S rRNA | 1136-1152 | 1136-1144, 1151-1152 | 37-53 | 37-45, 52-53 |
| A 18S rRNA | 1176-1200 | 1176-1186, 1196, 1199-1200 | 54-78 | 54-64, 74, 77-78 |
| A 18S rRNA | 1207-1210 | 1207-1210 | 79-82 | 79-82 |
| A 18S rRNA | 1263-1291 | 1263-1267, 1285-1291 | 83-111 | 83-87, 105-111 |
| A 18S rRNA | 1299-1302 | 1299-1302 | 112-115 | 112-115 |
| A 18S rRNA | 1327-1329 | 1327-1329 | 116-118 | 116-118 |
| A 18S rRNA | 1419-1449 | 1419-1421, 1442-1449 | 119-149 | 119-121, 142-149 |
| A 18S rRNA | 1461-1465 | 1461-1465 | 150-154 | 150-154 |
| A 18S rRNA | 1621-1647 | 1621-1629, 1642-1647 | 155-181 | 155-163, 176-181 |
| A 18S rRNA | 1752-1772 | 1752-1753, 1766, 1769-1772 | 182-202 | 182-183, 196, 199-202 |
| A 18S rRNA | 1780-1783 | 1780-1783 | 203-206 | 203-206 |
| AB S3 | 26-27 | 26-27 | 207-208 | 207-208 |
| AB S3 | 102-120 | 102-114, 117-120 | 209-227 | 209-221, 224-227 |
| AB S3 | 134-159 | 134-135, 153-159 | 228-253 | 228-229, 247-253 |
| AB S3 | 170-187 | 170-171, 186-187 | 254-271 | 254-255, 270-271 |
| AC S29 | 24-29 | 24-29 | 272-277 | 272-277 |
| AC S29 | 39-45 | 39-45 | 278-284 | 278-284 |
| B 25S rRNA | 2253-2264 | 2253-2255, 2258-2264 | 285-296 | 285-287, 290-296 |
| BC S30 | 2-17 | 2-17 | 297-312 | 297-312 |
| CC S31 | 82-85 | 82-85 | 313-316 | 313-316 |
| DC eEF2 | 574-588 | 574-576, 588 | 317-331 | 317-319, 331 |
| DC eEF2 | 608-612 | 608-612 | 332-336 | 332-336 |
| DC eEF2 | 631-635 | 631-635 | 337-341 | 337-341 |
| DC eEF2 | 646-669 | 646-649, 667-669 | 342-365 | 342-345, 363-365 |
| DC eEF2 | 686-713 | 686-689, 710-713 | 366-393 | 366-369, 390-393 |
| DC eEF2 | 839-842 | 839-842 | 394-397 | 394-397 |
| EC tRNA (IRES) | 6895-6915 | 6895-6897, 6913-6915 | 398-418 | 398-400, 416-418 |
| EC mRNA (IRES) | 6946-6958 | 6946, 6958 | 419-431 | 419, 431 |
| HB S10 | 59-62 | 59-62 | 432-435 | 432-435 |
| NB S16 | 137-143 | 137-143 | 436-442 | 436-442 |
| RB S20 | 67-82 | 67, 77-82 | 443-458 | 443, 453-458 |
| UB S23 | 58-70 | 58-59, 68-70 | 459-471 | 459-460, 469-471 |
| UB S23 | 115-118 | 115-118 | 472-475 | 472-475 |

| | | | | |
|---|---|---|---|---|
| **ZA S2** | 83-98 | 83-86, 97-98 | 476-491 | 476-479, 490-491 |
| **ZA S2** | 118-120 | 118-120 | 492-494 | 492-494 |

*Same residue numbering for 5JUO, 5JUS, 5JUT, and 5JUU.

#### *1.3 Supplementary Videos*

**Video S1.** Trajectory with unmethylated R146 (separate file).
Representative trajectory sampled at 2 frames per ns (R146 trajectory number 16; Table S1A).

**Video S2.** Trajectory with methylated R146 (separate file).
Representative trajectory sampled at 2 frames per ns (mR146 trajectory number 1; Table S1B).

#### *1.4 Supplementary Data File*

**Data File S1**. Computer code for molecular dynamics analysis (separate file).
