## Supplemental Data File S1 for "Arginine Methylation Regulates Ribosome CAR Function"

```
DYNAMICS

Energy Minimization
Heating (20ps)
Equilibration (3ns)
Neutral Dynamics (60ns, 100ns)

ANALYSIS

RMSD - cpptraj 1
COMdist - cpptraj 3
RMSF - cpptraj 4
avgHbond - cpptraj 5
RMS2D - cpptraj 7
SASA - cpptraj 9

SYNTAX

Energy Minimization

Energy Minimization Input Scripts:

emin1.in

# 20000 steps of minimization with explicit solvent and ions and 100.0 kcal/mol-A restraints on protein.
 &cntrl
 	maxcyc=20000,    ! number of cycles of minimization
     imin=1,    	 ! energy minimization on
     ntmin=1,    	 ! switch from steepest descent to conjugate gradient
     ncyc=2500,     ! switch method after 2500 cycles
     cut=9.0,    	 ! non-bonded cutoff distance
     igb=0,    	 ! solvent model
     ntb=1,    	 ! constant volume periodic boundaries
     ntpr=10,    	 ! report output every 10 steps
     ntr=1,   		 ! restraint on
 	restraint_wt=100.0,
     restraintmask=':1-494',
 &end

emin2.in

# 10000 steps of minimization with explicit solvent and ions and 75.0 kcal/mol-A restraints on protein.
 &cntrl
 	maxcyc=10000,    ! number of cycles of minimization
     imin=1,    	 ! energy minimization on
     ntmin=1,    	 ! switch from steepest descent to conjugate gradient
     ncyc=2500,     ! switch method after 2500 cycles
     cut=9.0,    	 ! non-bonded cutoff distance
     igb=0,    	 ! solvent model
     ntb=1,    	 ! constant volume periodic boundaries
     ntpr=10,    	 ! report output every 10 steps
     ntr=1,   		 ! restraint on
 	restraint_wt=75.0,
     restraintmask=':1-494',
 &end

emin3.in

# 5000 steps of minimization with explicit solvent and ions and 65.0 kcal/mol-A restraints on protein.
 &cntrl
 	maxcyc=5000,    ! number of cycles of minimization
     imin=1,    	 ! energy minimization on
     ntmin=1,    	 ! switch from steepest descent to conjugate gradient
     ncyc=2500,     ! switch method after 2500 cycles
     cut=9.0,    	 ! non-bonded cutoff distance
     igb=0,    	 ! solvent model
     ntb=1,    	 ! constant volume periodic boundaries
     ntpr=10,    	 ! report output every 10 steps
     ntr=1,   		 ! restraint on
 	restraint_wt=65.0,
     restraintmask=':1-494',
 &end

emin4.in

# 3000 steps of minimization with explicit solvent and ions and 55.0 kcal/mol-A restraints on protein.
 &cntrl
 	maxcyc=3000,    ! number of cycles of minimization
     imin=1,    	 ! energy minimization on
     ntmin=1,    	 ! switch from steepest descent to conjugate gradient
     ncyc=2500,     ! switch method after 2500 cycles
     cut=9.0,    	 ! non-bonded cutoff distance
     igb=0,    	 ! solvent model
     ntb=1,    	 ! constant volume periodic boundaries
     ntpr=10,    	 ! report output every 10 steps
     ntr=1,   		 ! restraint on
 	restraint_wt=55.0,
     restraintmask=':1-494',
 &end

emin5.in

# 3000 steps of minimization with explicit solvent and ions and 45.0 kcal/mol-A restraints on protein.
 &cntrl
 	maxcyc=3000,    ! number of cycles of minimization
     imin=1,    	 ! energy minimization on
     ntmin=1,    	 ! switch from steepest descent to conjugate gradient
     ncyc=2500,     ! switch method after 2500 cycles
     cut=9.0,    	 ! non-bonded cutoff distance
     igb=0,    	 ! solvent model
     ntb=1,    	 ! constant volume periodic boundaries
     ntpr=10,    	 ! report output every 10 steps
     ntr=1,   		 ! restraint on
 	restraint_wt=45.0,
     restraintmask=':1-494',
 &end

emin6.in

# 2000 steps of minimization with explicit solvent and ions and 30.0 kcal/mol-A restraints on protein.
 &cntrl
 	maxcyc=2000,    ! number of cycles of minimization
     imin=1,    	 ! energy minimization on
     ntmin=1,    	 ! switch from steepest descent to conjugate gradient
     ncyc=2000,     ! switch method after 2000 cycles
     cut=9.0,    	 ! non-bonded cutoff distance
     igb=0,    	 ! solvent model
     ntb=1,    	 ! constant volume periodic boundaries
     ntpr=10,    	 ! report output every 10 steps
     ntr=1,   		 ! restraint on
 	restraint_wt=30.0,
     restraintmask=':1-494',
 &end

emin7.in

# 2000 steps of minimization with explicit solvent and ions and 20.0 kcal/mol-A restraints on protein.
 &cntrl
 	maxcyc=2000,    ! number of cycles of minimization
     imin=1,    	 ! energy minimization on
     ntmin=1,    	 ! switch from steepest descent to conjugate gradient
     ncyc=2000,     ! switch method after 2000 cycles
     cut=9.0,    	 ! non-bonded cutoff distance
     igb=0,    	 ! solvent model
     ntb=1,    	 ! constant volume periodic boundaries
     ntpr=10,    	 ! report output every 10 steps
     ntr=1,   		 ! restraint on
 	restraint_wt=20.0,
     restraintmask=':1-494',
 &end

emin8.in

# 2000 steps of minimization with explicit solvent and ions and 15.0 kcal/mol-A restraints on protein.
 &cntrl
 	maxcyc=2000,    ! number of cycles of minimization
     imin=1,    	 ! energy minimization on
     ntmin=1,    	 ! switch from steepest descent to conjugate gradient
     ncyc=2000,     ! switch method after 2000 cycles
     cut=9.0,    	 ! non-bonded cutoff distance
     igb=0,    	 ! solvent model
     ntb=1,    	 ! constant volume periodic boundaries
     ntpr=10,    	 ! report output every 10 steps
     ntr=1,   		 ! restraint on
 	restraint_wt=15.0,
     restraintmask=':1-494',
 &end

emin9.in

# 2000 steps of minimization with explicit solvent and ions and 10.0 kcal/mol-A restraints on protein.
 &cntrl
 	maxcyc=2000,    ! number of cycles of minimization
     imin=1,    	 ! energy minimization on
     ntmin=1,    	 ! switch from steepest descent to conjugate gradient
     ncyc=2000,     ! switch method after 2000 cycles
     cut=9.0,    	 ! non-bonded cutoff distance
     igb=0,    	 ! solvent model
     ntb=1,    	 ! constant volume periodic boundaries
     ntpr=10,    	 ! report output every 10 steps
     ntr=1,   		 ! restraint on
 	restraint_wt=10.0,
     restraintmask=':1-494',
 &end

emin10.in

# 2000 steps of minimization with explicit solvent and ions and 5.0 kcal/mol-A restraints on protein.
 &cntrl
 	maxcyc=2000,    ! number of cycles of minimization
     imin=1,    	 ! energy minimization on
     ntmin=1,    	 ! switch from steepest descent to conjugate gradient
     ncyc=2000,     ! switch method after 2000 cycles
     cut=9.0,    	 ! non-bonded cutoff distance
     igb=0,    	 ! solvent model
     ntb=1,    	 ! constant volume periodic boundaries
     ntpr=10,    	 ! report output every 10 steps
     ntr=1,   		 ! restraint on
 	restraint_wt=5.0,
     restraintmask=':1-494',
 &end

emin11.in

# 2000 steps of minimization with explicit solvent and ions and 1.0 kcal/mol-A restraints on protein.
 &cntrl
 	maxcyc=2000,    ! number of cycles of minimization
     imin=1,    	 ! energy minimization on
     ntmin=1,    	 ! switch from steepest descent to conjugate gradient
     ncyc=2000,     ! switch method after 2500 cycles
     cut=9.0,    	 ! non-bonded cutoff distance
     igb=0,    	 ! solvent model
     ntb=1,    	 ! constant volume periodic boundaries
     ntpr=10,    	 ! report output every 10 steps
     ntr=1,   		 ! restraint on
 	restraint_wt=1.0,
     restraintmask=':1-494',
 &end


Heating (20ps)

Heating Input Script:

20ps_heat.in

heating: MD with restraint on molecule
 &cntrl
  imin=0,   	 ! no minimization
  irest=0,   	 ! randomly assign velocities
  ntx=1,   	 ! randomly assign velocities
  ntb=1,   	 ! periodic boundaries for constant volume
  cut=10,   	 ! non-bond cutoff of 10 angstroms
  ntr=1,    	 ! restraints on
  ntc=2,    	 ! SHAKE on
  ntf=2,   	 ! bond interactions involving H omitted
  tempi=0.0,    ! initial temperature
  temp0=300.0,    ! ref temperature
  ntt=3,    	 ! Langevin dynamics
  gamma_ln=1.0,    ! collision frequency
  nstlim=10000, ! number of MD steps to be performed (20ps)
  dt=0.002,   	 ! timestep in ps
  ntpr=1000,     ! print energy every 'ntpr' steps
  ntwx=1000,     ! write coord to trj every 'ntwx' steps
  ntwr=10000,    ! rewrite rst file every 'ntwr' steps
  restraint_wt=20.0,
  restraintmask=':1 - 494'
 /

Equilibration (3ns)

Equilibration Input Script:

3ns_equil.in

3ns equilibration step (EQUIL)
 &cntrl
  imin 	= 0,   		 ! no minimization
  ntx  	= 5,   		 ! velocities inherited
  irest	= 1,   		 ! velocities inherited
  ntpr 	= 5000,   	 ! print energy info every `ntpr` steps
  ntwr 	= 50000,   	 ! rewrite rst file every `ntwr` steps
  ntwx 	= 1000,   	 ! write coord to trj every `ntwx` steps
  ntf  	= 2,   		 ! bond interactions involving H omitted
  ntc  	= 2,   		 ! SHAKE on, Hbonds constrained
  cut  	= 8.0,   	 ! non-bond cutoff of 8A
  ntb  	= 2,   		 ! periodic boundaries for constant pressure
  nstlim   = 3000000,    ! number of MD steps to be performed (3ns)
  dt   	= 0.001,   	 ! time step in psec
  tempi	= 0.0,   	 ! initial temperature
  temp0	= 300,   	 ! ref temperature
  ntt  	= 3,   		 ! Langevin dynamics
  gamma_ln = 1.0,   	 ! collision freq
  ntp  	= 1,   		 ! constant pressure dynamics
  pres0	= 1.0,   	 ! reference pressure 1
  taup 	= 5.0,   	 ! time constant for pressure
  nmropt   = 1,   		 ! restraint on
  ioutfm   = 1,   		 ! write binary trajectory
  ntr  	= 1,   		 ! restraint on
  restraint_wt = 20.0,
  restraintmask=':1-494',
  /
 &end
 &wt
  type='END',
 &end


Neutral Dynamics (60ns, 100ns)

Neutral Dynamics Input Scripts:

60ns_nd.in

60 ns neutral dynamics (NEUTRAL)
 &cntrl
  imin 	= 0,   		 ! no minimization
  ntx  	= 5,   		 ! velocities inherited
  irest	= 1,   		 ! velocities inherited
  ntpr 	= 5000,    	 ! print energy info every `ntpr` steps
  ntwr 	= 50000,   	 ! rewrite rst file every `ntwr` steps
  ntwx 	= 5000,    	 ! write coord to trj every `ntwx` steps
  ntf  	= 2,   		 ! bond interactions involving H omitted
  ntc  	= 2,   		 ! SHAKE on, Hbonds constrained
  cut  	= 8.0, 		 ! non-bond cutoff of 8A
  ntb  	= 2,   		 ! 2 periodic boundaries for constant pressure
  nstlim   = 30000000,    ! number of MD steps to be performed (60ns)
  dt   	= 0.002,   	 ! time step in psec
  tempi	= 0.0, 		 ! initial temperature
  temp0	= 300, 		 ! ref temperature
  ntt  	= 3,   		 ! Langevin dynamics
  gamma_ln = 1.0, 		 ! collision freq
  ntp  	= 1,   		 ! constant pressure dynamics
  pres0	= 1.0, 		 ! reference pressure 1
  taup 	= 5.0, 		 ! time constant for pressure
  nmropt   = 1,   		 ! restraint on
  ioutfm   = 1,   		 ! write binary trajectory
  ntr  	= 1,   		 ! restraint on
  restraint_wt = 20.0,
  restraintmask=':1,6-12,17-22,32-45,52-64,74,77-87,105-121,142-163,176-183,196,199-221,224-229,247-255,270-287,290-319,331-345,363-369,390-400,416-419,431-443,453-460,469-479,490-494',
  /
 &end
 &wt
  type='END',
 &end

100ns_nd.in

100 ns neutral dynamics (NEUTRAL)
 &cntrl
  imin 	= 0,   		 ! no minimization
  ntx  	= 5,   		 ! velocities inherited
  irest	= 1,   		 ! velocities inherited
  ntpr 	= 5000,    	 ! print energy info every `ntpr` steps
  ntwr 	= 50000,   	 ! rewrite rst file every `ntwr` steps
  ntwx 	= 5000,    	 ! write coord to trj every `ntwx` steps
  ntf  	= 2,   		 ! bond interactions involving H omitted
  ntc  	= 2,   		 ! SHAKE on, Hbonds constrained
  cut  	= 8.0, 		 ! non-bond cutoff of 8A
  ntb  	= 2,   		 ! 2 periodic boundaries for constant pressure
  nstlim   = 50000000,    ! number of MD steps to be performed (100ns)
  dt   	= 0.002,   	 ! time step in psec
  tempi	= 0.0, 		 ! initial temperature
  temp0	= 300, 		 ! ref temperature
  ntt  	= 3,   		 ! Langevin dynamics
  gamma_ln = 1.0, 		 ! collision freq
  ntp  	= 1,   		 ! constant pressure dynamics
  pres0	= 1.0, 		 ! reference pressure 1
  taup 	= 5.0, 		 ! time constant for pressure
  nmropt   = 1,   		 ! restraint on
  ioutfm   = 1,   		 ! write binary trajectory
  ntr  	= 1,   		 ! restraint on
  restraint_wt = 20.0,
  restraintmask=':1,6-12,17-22,32-45,52-64,74,77-87,105-121,142-163,176-183,196,199-221,224-229,247-255,270-287,290-319,331-345,363-369,390-400,416-419,431-443,453-460,469-479,490-494',
  /
 &end
 &wt
  type='END',
 &end

RMSD - cpptraj 1

Strip Trajectory:

# This script is to strip the trajectories.
# prmtop file
parm /mindstore/home33ext/kscopino/mR146_N2/5JUP/GCU/NO_MOD/TLEAP/5JUP_N2_NM_mR146_wat.prmtop [modi3]

# experimental neutral dynamics trajectory
trajin /mindstore/home33ext/kscopino/mR146_N2/5JUP/GCU/NO_MOD/NEUTRAL/NEUTRAL_10/mdcrd_nd_10 parm [modi3]

autoimage
strip :WAT
strip :K+
trajout ../mdcrd_nd_10_strip nobox

Average Structure:

# This script is to create an average structure for each interval of a trajectory.

# prmtop file
parm /mindstore/home33ext/kscopino/mR146_N2/5JUP/GCU/NO_MOD/TLEAP/5JUP_N2_NM_mR146_nowat.prmtop

# experimental neutral dynamics trajectory
trajin /mindstore/home33ext/kscopino/mR146_N2/5JUP/GCU/NO_MOD/NEUTRAL/NEUTRAL_20/mdcrd_nd_20_strip

# make average
average avg_struct.rst restart

RMSD Calculation:

Average Reference Version:

# This script is to check the stability of an experimental run using RMSD to determine the length of equilibration dynamics
# prmtop file
parm /mindstore/home33ext/kscopino/mR146_N2/5JUP/GCU/NO_MOD/TLEAP/5JUP_N2_NM_mR146_nowat.prmtop [modi3]

# reference for RMSD (should be the average structure for the entire trajectory)
trajin /mindstore/home33ext/kscopino/mR146_N2/5JUP/GCU/NO_MOD/NEUTRAL/NEUTRAL_24/DATA/avg_struct.rst

# experimental neutral dynamics trajectory
trajin /mindstore/home33ext/kscopino/mR146_N2/5JUP/GCU/NO_MOD/NEUTRAL/NEUTRAL_24/mdcrd_nd_24_strip parm [modi3]

# RMSD of unrestrained backbone atoms
rms unrestr_resid_A first :2-5,13-16,23-31,46-51,65-73,75-76,88-104,122-141,164-175,184-195,197-198,222-223,230-246,256-269,288-289,320-330,346-362,370-389,401-415,420-430,444-452,461-468,480-489@N,CA,C,O,P,O5',O3',C5' out rmsd_24_NOTrestr_bkbone_A.dat
# RMSD of restrained backbone atoms
rms restr_resid_A first :1,6-12,17-22,32-45,52-64,74,77-87,105-121,142-163,176-183,196,199-221,224-229,247-255,270-287,290-319,331-345,363-369,390-400,416-419,431-443,453-460,469-479,490-494@N,CA,C,O,P,O5',O3',C5' out rmsd_24_restr_bkbone_A.dat
# RMSD of mRNA (A-site codon, +1 codon), tRNA anticodon, and CAR
rms unrestr_resid_local_A first :94,127,240,408-410,423-428@N,CA,C,O,P,O5',O3',C5' out rmsd_24_local_bkbone_A.dat

Equilibration Out Reference Version:

# This script is to check the stability of an experimental run using RMSD to determine the length of equilibration dynamics
# prmtop file
parm /mindstore/home33ext/kscopino/mR146_N2/5JUP/GCU/NO_MOD/TLEAP/5JUP_N2_NM_mR146_wat.prmtop [modi3]

# reference for RMSD (should be the equilibration out structure)
trajin /mindstore/home33ext/kscopino/mR146_N2/5JUP/GCU/NO_MOD/EQUIL/EQUIL_24/5JUP_N2_NM_mR146_equil_24.rst

# experimental neutral dynamics trajectory
trajin /mindstore/home33ext/kscopino/mR146_N2/5JUP/GCU/NO_MOD/NEUTRAL/NEUTRAL_24/mdcrd_nd_24 parm [modi3]

# RMSD of unrestrained backbone atoms
rms unrestr_resid_EO first :2-5,13-16,23-31,46-51,65-73,75-76,88-104,122-141,164-175,184-195,197-198,222-223,230-246,256-269,288-289,320-330,346-362,370-389,401-415,420-430,444-452,461-468,480-489@N,CA,C,O,P,O5',O3',C5' out rmsd_24_NOTrestr_bkbone_EO.dat
# RMSD of restrained backbone atoms
rms restr_resid_EO first :1,6-12,17-22,32-45,52-64,74,77-87,105-121,142-163,176-183,196,199-221,224-229,247-255,270-287,290-319,331-345,363-369,390-400,416-419,431-443,453-460,469-479,490-494@N,CA,C,O,P,O5',O3',C5' out rmsd_24_restr_bkbone_EO.dat
# RMSD of mRNA (A-site codon, +1 codon), tRNA anticodon, and CAR
rms unrestr_resid_local_EO first :94,127,240,408-410,423-428@N,CA,C,O,P,O5',O3',C5' out rmsd_24_local_bkbone_EO.dat


COMdist - cpptraj 3

COMdist Calculation:

# This script is to examine the stacking of the CAR residues.
# It looks at the COM distance between residues participating in pi-stacking.

# prmtop file
parm /mindstore/home33ext/kscopino/COD1_SUBS_N2/5JUP/GCU/NO_MOD/TLEAP/5JUP_N2_1_GC_wat.prmtop [modi3]

# experimental neutral dynamics trajectory
trajin /mindstore/home33ext/kscopino/COD1_SUBS_N2/5JUP/GCU/NO_MOD/NEUTRAL/NEUTRAL_25/mdcrd_nd_25 parm [modi3]

# STACKING DISTANCES (COM)
# Base Stacking of nt35 (tRNA nt2, G) with nt34 (tRNA wobble, G)
distance d409C24568N1379_408C24568N1379 :409@C2,C4,C5,C6,C8,N1,N3,N7,N9 :408@C2,C4,C5,C6,C8,N1,N3,N7,N9 out dist_409_C24568N1379_408_C24568N1379_2to2ring.dat

# Base Stacking of nt34 (tRNA wobble, G) with C1054
distance d408C24568N1379_94C2456N13 :408@C2,C4,C5,C6,C8,N1,N3,N7,N9 :94@C2,C4,C5,C6,N1,N3 out dist_408_C24568N1379_94_C2456N13_2to1ring.dat

# Base Stacking of C1054 with A1196
distance d94C2456N13_127C24568N1379 :94@C2,C4,C5,C6,N1,N3 :127@C2,C4,C5,C6,C8,N1,N3,N7,N9 out dist_94_C2456N13_127_C24568N1379_1to2ring.dat

# Base Stacking of A1196 with R146
distance d127C24568N1379_R240CzNeNh1Nh2 :127@C2,C4,C5,C6,C8,N1,N3,N7,N9 :240@CZ,NE,NH1,NH2  out dist_127_C24568N1379_R240CzNeNh1Nh2_2toGUANring.dat


RMSF - cpptraj 4

RMSF Calculation:

# prmtop input
parm /mindstore/home33ext/kscopino/COD1_SUBS_N2/5JUP/GUU/NO_MOD/TLEAP/5JUP_N2_NM_GUU_nowat.prmtop

# trajectory to analyze
trajin /mindstore/home33ext/kscopino/COD1_SUBS_N2/5JUP/GUU/NO_MOD/NEUTRAL/NEUTRAL_28/mdcrd_nd_28_strip 2000

# RMS fit to average structure
average crdset avg_struct.rst
run
rms ref avg_struct.rst

# RMSF
# nt CU core atoms
rmsf out rmsf_ntCU_core.dat :94,423-425,427,428@C2,C4,C5,C6,N1,N3 byres
# nt AG core atoms
rmsf out rmsf_ntAG_core.dat :127,408-410,426@C2,C4,C5,C6,C8,N1,N3,N7,N9 byres
# aa R core atoms
rmsf out rmsf_aaR_core.dat :240@CZ,NE,NH1,NH2 byres


avgHbond - cpptraj 5

avgHbond Calculation:

parm /mindstore/home33ext/kscopino/mR146_N2/5JUP/GCU/NO_MOD/TLEAP/5JUP_N2_NM_mR146_wat.prmtop [modi3]
trajin /mindstore/home33ext/kscopino/mR146_N2/5JUP/GCU/NO_MOD/NEUTRAL/NEUTRAL_27/mdcrd_nd_27 2000 parm [modi3]
autoimage

### In-Registration ###
# Anticodon position 1/codon position 1
hbond nhb_AVE_410_all_423_all :410|:423 avgout nhb_AVE_410_all_423_all.dat
# Anticodon position 2/codon position 2
hbond nhb_AVE_409_all_424_all :409|:424 avgout nhb_AVE_409_all_424_all.dat
# Anticodon position 3/codon position 3
hbond nhb_AVE_408_all_425_all :408|:425 avgout nhb_AVE_408_all_425_all.dat
# C1054 to +1 N1
hbond nhb_AVE_94_all_426_all :94|:426 avgout nhb_AVE_94_all_426_all.dat
# A1196 to +1 N2
hbond nhb_AVE_127_all_427_all :127|:427 avgout nhb_AVE_127_all_427_all.dat
# R146 to +1 N2
hbond nhb_AVE_240_all_427_all :240|:427 avgout nhb_AVE_240_all_427_all.dat

### Cross-Registration ###
# Anticodon position 1/codon position 2
hbond nhb_AVE_410_all_424_all :410|:424 avgout nhb_AVE_410_all_424_all.dat
# Anticodon position 2/codon position 1
hbond nhb_AVE_409_all_423_all :409|:423 avgout nhb_AVE_409_all_423_all.dat
# Anticodon position 2/codon position 3
hbond nhb_AVE_409_all_425_all :409|:425 avgout nhb_AVE_409_all_425_all.dat
# Anticodon position 3/codon position 2
hbond nhb_AVE_408_all_424_all :408|:424 avgout nhb_AVE_408_all_424_all.dat
# Anticodon position 3 to N1
hbond nhb_AVE_408_all_426_all :408|:426 avgout nhb_AVE_408_all_426_all.dat
# C1054 to codon position 3
hbond nhb_AVE_94_all_425_all :94|:425 avgout nhb_AVE_94_all_425_all.dat
# C1054 to +1 N2
hbond nhb_AVE_94_all_427_all :94|:427 avgout nhb_AVE_94_all_427_all.dat
# C1054 to +1 N3
hbond nhb_AVE_94_all_428_all :94|:428 avgout nhb_AVE_94_all_428_all.dat
# A1196 to +1 N1
hbond nhb_AVE_127_all_426_all :127|:426 avgout nhb_AVE_127_all_426_all.dat
# A1196 to +1 N3
hbond nhb_AVE_127_all_428_all :127|:428 avgout nhb_AVE_127_all_428_all.dat
# R146 to +1 N1
hbond nhb_AVE_240_all_426_all :240|:426 avgout nhb_AVE_240_all_426_all.dat
# R146 to +1 N3
hbond nhb_AVE_240_all_428_all :240|:428 avgout nhb_AVE_240_all_428_all.dat


RMS2D - cpptraj 7

12-residue version:

# prmtop input
parm /mindstore/home33ext/kscopino/mR146_N2/5JUP/GCU/NO_MOD/TLEAP/5JUP_N2_NM_mR146_nowat.prmtop [modi3]

# trajectory to analyze, sampling 1 in 100 frames
trajin /mindstore/home33ext/kscopino/mR146_N2/5JUP/GCU/NO_MOD/NEUTRAL/NEUTRAL_21/mdcrd_nd_21_strip 2000 last 100 parm [modi3]
trajin /mindstore/home33ext/kscopino/mR146_N2/5JUP/GCU/NO_MOD/NEUTRAL/NEUTRAL_22/mdcrd_nd_22_strip 2000 last 100 parm [modi3]
trajin /mindstore/home33ext/kscopino/mR146_N2/5JUP/GCU/NO_MOD/NEUTRAL/NEUTRAL_23/mdcrd_nd_23_strip 2000 last 100 parm [modi3]
trajin /mindstore/home33ext/kscopino/mR146_N2/5JUP/GCU/NO_MOD/NEUTRAL/NEUTRAL_24/mdcrd_nd_24_strip 2000 last 100 parm [modi3]
trajin /mindstore/home33ext/kscopino/mR146_N2/5JUP/GCU/NO_MOD/NEUTRAL/NEUTRAL_25/mdcrd_nd_25_strip 2000 last 100 parm [modi3]
trajin /mindstore/home33ext/kscopino/mR146_N2/5JUP/GCU/NO_MOD/NEUTRAL/NEUTRAL_26/mdcrd_nd_26_strip 2000 last 100 parm [modi3]
trajin /mindstore/home33ext/kscopino/mR146_N2/5JUP/GCU/NO_MOD/NEUTRAL/NEUTRAL_27/mdcrd_nd_27_strip 2000 last 100 parm [modi3]
trajin /mindstore/home33ext/kscopino/mR146_N2/5JUP/GCU/NO_MOD/NEUTRAL/NEUTRAL_28/mdcrd_nd_28_strip 2000 last 100 parm [modi3]
trajin /mindstore/home33ext/kscopino/mR146_N2/5JUP/GCU/NO_MOD/NEUTRAL/NEUTRAL_29/mdcrd_nd_29_strip 2000 last 100 parm [modi3]
trajin /mindstore/home33ext/kscopino/mR146_N2/5JUP/GCU/NO_MOD/NEUTRAL/NEUTRAL_30/mdcrd_nd_30_strip 2000 last 100 parm [modi3]

autoimage

# calculate core atoms rms2d using only the A-site and +1 codon mRNA, tRNA, and CAR
rms2d 12res_coreatoms :94,127,240,408-410,423-428@C2,C4,C5,C6,C8,N1,N3,N7,N9,NH1,NH2,CZ,NE out rms2d_concat_12res_1in100_coreatoms_100ns.dat

# calculate backbone rms2d using only the A-site and +1 codon mRNA, tRNA, and CAR
rms2d 12res_bkbone :94,127,240,408-410,423-428@N,C,CA,O,P,O5',O3',C5' out rms2d_concat_12res_1in100_100ns.dat

11-residue version:

# prmtop input
parm /mindstore/home33ext/kscopino/mR146_N2/5JUP/GCU/NO_MOD/TLEAP/5JUP_N2_NM_mR146_nowat.prmtop [modi3]

# trajectory to analyze, sampling 1 in 100 frames
trajin /mindstore/home33ext/kscopino/mR146_N2/5JUP/GCU/NO_MOD/NEUTRAL/NEUTRAL_21/mdcrd_nd_21_strip 2000 last 100 parm [modi3]
trajin /mindstore/home33ext/kscopino/mR146_N2/5JUP/GCU/NO_MOD/NEUTRAL/NEUTRAL_22/mdcrd_nd_22_strip 2000 last 100 parm [modi3]
trajin /mindstore/home33ext/kscopino/mR146_N2/5JUP/GCU/NO_MOD/NEUTRAL/NEUTRAL_23/mdcrd_nd_23_strip 2000 last 100 parm [modi3]
trajin /mindstore/home33ext/kscopino/mR146_N2/5JUP/GCU/NO_MOD/NEUTRAL/NEUTRAL_24/mdcrd_nd_24_strip 2000 last 100 parm [modi3]
trajin /mindstore/home33ext/kscopino/mR146_N2/5JUP/GCU/NO_MOD/NEUTRAL/NEUTRAL_25/mdcrd_nd_25_strip 2000 last 100 parm [modi3]
trajin /mindstore/home33ext/kscopino/mR146_N2/5JUP/GCU/NO_MOD/NEUTRAL/NEUTRAL_26/mdcrd_nd_26_strip 2000 last 100 parm [modi3]
trajin /mindstore/home33ext/kscopino/mR146_N2/5JUP/GCU/NO_MOD/NEUTRAL/NEUTRAL_27/mdcrd_nd_27_strip 2000 last 100 parm [modi3]
trajin /mindstore/home33ext/kscopino/mR146_N2/5JUP/GCU/NO_MOD/NEUTRAL/NEUTRAL_28/mdcrd_nd_28_strip 2000 last 100 parm [modi3]
trajin /mindstore/home33ext/kscopino/mR146_N2/5JUP/GCU/NO_MOD/NEUTRAL/NEUTRAL_29/mdcrd_nd_29_strip 2000 last 100 parm [modi3]
trajin /mindstore/home33ext/kscopino/mR146_N2/5JUP/GCU/NO_MOD/NEUTRAL/NEUTRAL_30/mdcrd_nd_30_strip 2000 last 100 parm [modi3]

autoimage

# calculate core atoms rms2d using only the A-site and +1 codon mRNA, tRNA, and CA
rms2d 11res_coreatoms :94,127,408-410,423-428@C2,C4,C5,C6,C8,N1,N3,N7,N9 out rms2d_concat_11res_1in100_coreatoms_100ns.dat

# calculate backbone rms2d using only the A-site and +1 codon mRNA, tRNA, and CA
rms2d 11res_bkbone :94,127,408-410,423-428@N,C,CA,O,P,O5',O3',C5' out rms2d_concat_11res_1in100_100ns.dat

SASA – cpptraj 9

Strip Trajectories:

# This script is to strip the trajectories for SASA.
# prmtop file
parm /mindstore/home33ext/kscopino/COD1_SUBS_N2/5JUP/GCU/NO_MOD/TLEAP/5JUP_N2_AR.prmtop [modi3]

# experimental neutral dynamics trajectory
#trajin /mindstore/home33ext/kscopino/mR146_N2/5JUP/GCU/NO_MOD/NEUTRAL/NEUTRAL_26/mdcrd_nd_26_strip parm [modi3]

trajin /mindstore/home33ext/kscopino/mR146_N2/5JUP/GCU/NO_MOD/NEUTRAL/NEUTRAL_26/mdcrd_nd_26_AR_strip parm [modi3]

autoimage
#strip :127@P,OP1,OP2,O5',C5',H5',H5'',C4',H4',O4',C1',H1',H8,H61,H62,H2,C3',H3',C2',H2',O2',HO2',O3'
#strip :240@C1,HC11,HC12,HH2,HNE,HCD1,HCD2,HCG1,HCG2,HCB1,HCB2,HCA,N,H,CA,CB,CG,CD,HH11,HH12,C,O
#strip :1-126,128-239,241-494

#strip :1

strip :2

trajout ../mdcrd_nd_26_A_strip nobox

SASA Calculation:

# This script is to examine the stacking between A and R of the CAR residues.
# prmtop file
parm /mindstore/home33ext/kscopino/mR146_N2/5JUP/GCU/NO_MOD/TLEAP/5JUP_N2_R.prmtop [modi3]

# experimental neutral dynamics trajectory
#trajin /mindstore/home33ext/kscopino/mR146_N2/5JUP/GCU/NO_MOD/NEUTRAL/NEUTRAL_27/mdcrd_nd_27_AR_strip parm [modi3]

#trajin /mindstore/home33ext/kscopino/mR146_N2/5JUP/GCU/NO_MOD/NEUTRAL/NEUTRAL_27/mdcrd_nd_27_A_strip parm [modi3]

trajin /mindstore/home33ext/kscopino/mR146_N2/5JUP/GCU/NO_MOD/NEUTRAL/NEUTRAL_27/mdcrd_nd_27_R_strip parm [modi3]

# SASA surrounding A1196 and R146
#surf surf_AR :1-2 out surf_AR.dat

# SASA surrounding A1196
#surf surf_A :1 out surf_A.dat

# SASA surrounding R146
surf surf_R :1 out surf_R.dat
```
